# Fine-Tuning Protein Language Models Enhances the Identification and Interpretation of the Transcription Factors

**DOI:** 10.1101/2025.11.27.691010

**Authors:** Mir Tanveerul Hassan, Saima Gaffar, Hamza Zahid, Sang Jun Lee

**Affiliations:** Jeonbuk RICE Intelligence Innovation Research Center, Jeonbuk National University, Jeonju, 54896, South Korea; Division of Electronics and Information Engineering, Jeonbuk National University, Jeonju, 54896, South Korea

**Keywords:** transcription factors, protein language model, full fine-tuning, parameter-efficient fine-tuning (PEFT), low-rank adaption (LoRA)

## Abstract

Transcription factors (TFs) are pivotal regulators of gene expression and play essential roles in diverse cellular activities. The three-dimensional organization of the genome and transcriptional regulation are predominantly orchestrated by TFs. By recruiting the transcriptional machinery to gene enhancers or promoters, TFs can either activate or repress transcription, thereby controlling gene activity and various biological pathways. Accurate identification of TFs is vital for elucidating gene regulatory mechanisms within cells. However, experimental identification remains labor-intensive and time-consuming, highlighting the necessity for efficient computational approaches. In this study, we present a two-layer predictive framework utilizing protein language models (pLMs) via full fine-tuning and parameter-efficient fine-tuning. The initial layer robustly classifies and identifies transcription factors, while the subsequent layer predicts TFs with a binding preference for methylated DNA (TFPMs). Our approach further incorporates attention weights and protein sequence motifs to enhance interpretability and predictive capability. By leveraging attention mechanisms, we highlight biologically relevant regions of the protein sequences that contribute most strongly to the predictions. Additionally, motif analysis facilitates the identification of conserved sequence patterns that are critical for TF recognition and function. Across both TF and TFPM classification tasks, the inclusion of these features allowed our methods to consistently surpass contemporary models, as demonstrated by independent test results.

**Keypoints:**

- Developed a two-layer predictive framework using protein language models (pLMs) with both full fine-tuning and parameter-efficient fine-tuning methods.
- The first layer accurately identifies transcription factors (TFs), and the second layer predicts TFs with binding preference for methylated DNA (TFPMs).
- Integrated attention weights and protein sequence motifs to enhance model interpretability by highlighting biologically relevant sequence regions and conserved patterns.
- Achieved superior performance compared to state-of-the-art methods, validated by independent testing.

**Mir Tanveerul Hassan** obtained his M.Tech. in Computer Science from the University of Kashmir, India, in 2020, and later earned his Ph.D. in Electronics and Information Engineering from Jeonbuk National University, Jeonju, South Korea. He is currently serving as a postdoctoral fellow at the Jeonbuk RICE Intelligence Innovation Research Center. His research interests encompass computational biology, bioinformatics, and pattern recognition.

**Saima Gaffar** received her B.Tech. and M.Tech. degrees in Computer Science from the University of Kashmir, Srinagar, India, and her Ph.D. in Electronics and Information Engineering from Jeonbuk National University, South Korea. Her research focuses on bioinformatics, computational biology, deep learning, and image processing.

**Hamza Zahid** received his B.S. degree in Mechatronics Engineering from the University of Engineering and Technology, Peshawar, Pakistan. He is currently pursuing the integrated M.S. and Ph.D. degrees in Electronics and Information Engineering at Jeonbuk National University, South Korea. His primary research interests include the applications of artificial intelligence in computational drug discovery.

**Sang Jun Lee** received his B.S., M.S., and Ph.D. degrees in Electrical Engineering from POSTECH, South Korea. Following his doctoral studies, he worked as a senior researcher at the Samsung Advanced Institute of Technology (SAIT). He is currently an Associate Professor in the Division of Electronics and Information Engineering at Jeonbuk National University, South Korea. His research interests include image analysis, deep learning, and medical image processing.

## Introduction

Transcription factors (TFs) are key regulatory proteins that play a central role in cell-fate determination by governing gene expression. They are distributed throughout the genome, and their abundance generally scales with genome size [1]. In fact, larger genomes tend to encode a greater number of TFs. It is estimated that approximately 10% of the human genome encodes TFs, making them one of the most prevalent protein families in humans. Acting as critical regulators of transcription, TFs interact with promoter regions to control gene expression by recruiting RNA polymerases to the core promoter, thereby either enhancing or repressing transcription. Although all cells share the same genetic blueprints, TFs drive cellular diversity by selectively modulating the expression of distinct gene sets, thus orchestrating processes such as differentiation. They regulate multiple target genes across distinct developmental stages, yet their activity is confined by specific DNA binding motifs. Mutations within TFs have been implicated in numerous diseases [2]. Essentially, TFs serve as molecular switches that govern the spatial, temporal, and quantitative dynamics of gene expression, ensuring precise transcriptional control. Accurate identification of TFs thus represents both a fundamental biological challenge and a major therapeutic opportunity for drug discovery. Structurally, TFs possess three primary functional domains: a DNA-binding domain, a transcriptional regulatory interface, and a signaling region [3, 4], which enable effective binding to gene promoters [5].

DNA methylation was initially recognized as a mechanism for influencing TF binding affinity, predominantly targeting cytosines within CpG dinucleotides [6]. Initially, methylation was regarded as a transcriptional repressive mark, especially when localized within CpG islands at promoters, where it obstructed transcription factor access and resulted in gene silencing. With advances in high-throughput and third-generation sequencing technologies, such as bisulfite sequencing, PacBio single-molecule real-time sequencing, and Oxford Nanopore sequencing, a much broader landscape of DNA modifications has been identified [7]. More recent studies further demonstrated that certain TFs, even in the absence of methyl-CpG binding domains (MBDs), retain the capacity to bind methylated DNA [8, 9, 10]. Elucidating these interactions is vital for understanding the biological diversity conferred by DNA methylation-dependent regulation [11, 12, 13].

Conventionally, TF discovery has relied on wet-lab approaches, including SELEX-based methods [14], MITOMI [15], and ChIP-based assays [16], which have enabled the identification of many TFs cataloged in public databases [17, 18, 19, 20]. However, these experimental strategies are resource-intensive in terms of both time and cost and remain inadequate for characterizing TFs across all tissues and species. Computational methods, particularly those driven by artificial intelligence, have emerged as powerful complements to experimental discovery. By exploiting models trained on known TF sequences, novel TF candidates can be computationally predicted and later validated through experiments, thereby streamlining the identification pipeline and dramatically reducing the associated costs.

Liu et al. (2020) [21] introduced TFPred, the first computational framework for distinguishing TFs from non-TFs, utilizing classical hand-crafted encodings such as composition/transition/distribution (CTD), amino acid composition (AAC), and dipeptide composition (DPC) with a support vector machine (SVM) classifier. However, these descriptors lacked the ability to capture sequential information, overlooking the positional arrangement of amino acids that can critically influence protein structure and function. To address this, Nguyen (2022) [22] proposed PSSM+CNN, wherein protein sequences were transformed into position-specific scoring matrices (PSSM) followed by classification with a deep convolutional neural network. Nonetheless, the accuracy of PSSM is contingent on sequence alignment quality, which introduces potential variability. Subsequently, Zheng et al. [23] adapted capsule networks (Capsule TF), inspired by Sabour et al. [24], while Li et al. (2022) [25] developed Li RNN, which directly processed protein sequences as residue triplets through long short-term memory (LSTM) networks. The latest model is the TFProtBert [26], using ProtBert embeddings as an input for the machine learning models, as the embedding vectors capture general semantic similarity but not task-specific distinctions. Despite their innovations, these neural models often suffered from overfitting due to limited training data and the complexity of tuning numerous hyperparameters.

Recent progress in transformer architectures and large language models (LLMs) has revolutionized protein sequence analysis. By drawing parallels between proteins and natural language, LLMs can model both local and long-range dependencies among amino acids, uncovering complex structural and functional relationships. This paradigm treats amino acid sequences as linguistic tokens, enabling pretrained models to capture rich contextual information that traditional encodings often overlook. This study uses ESM2 and ProtBert models for protein analysis. ESM2, trained on large protein datasets like UniRef50 via masked language modeling, captures rich biochemical and structural context without needing evolutionary alignments, aiding tasks like structure and function prediction. ProtBert, based on BERT, is pretrained on 217 million sequences from UniRef100, masking 15% of amino acids, and treats each protein as an independent document without next sentence prediction.

The full model fine-tuning and Parameter-Efficient FineTuning (PEFT) pLMs can achieve and show better results due to the inherent nature of large language models (LLMs) to capture context and subtle biological dependencies within protein sequences. Building on this insight, we propose leveraging pLMs with both full and parameter-efficient fine-tuning strategies to overcome the challenges of traditional methods. This approach demonstrates superior effectiveness in transcription factor (TF) classification and enables accurate prediction of TF interactions with methylated DNA, providing a powerful tool for understanding gene regulation mechanisms. The architecture of the proposed method is depicted in Figure 1.

**Figure 1.**
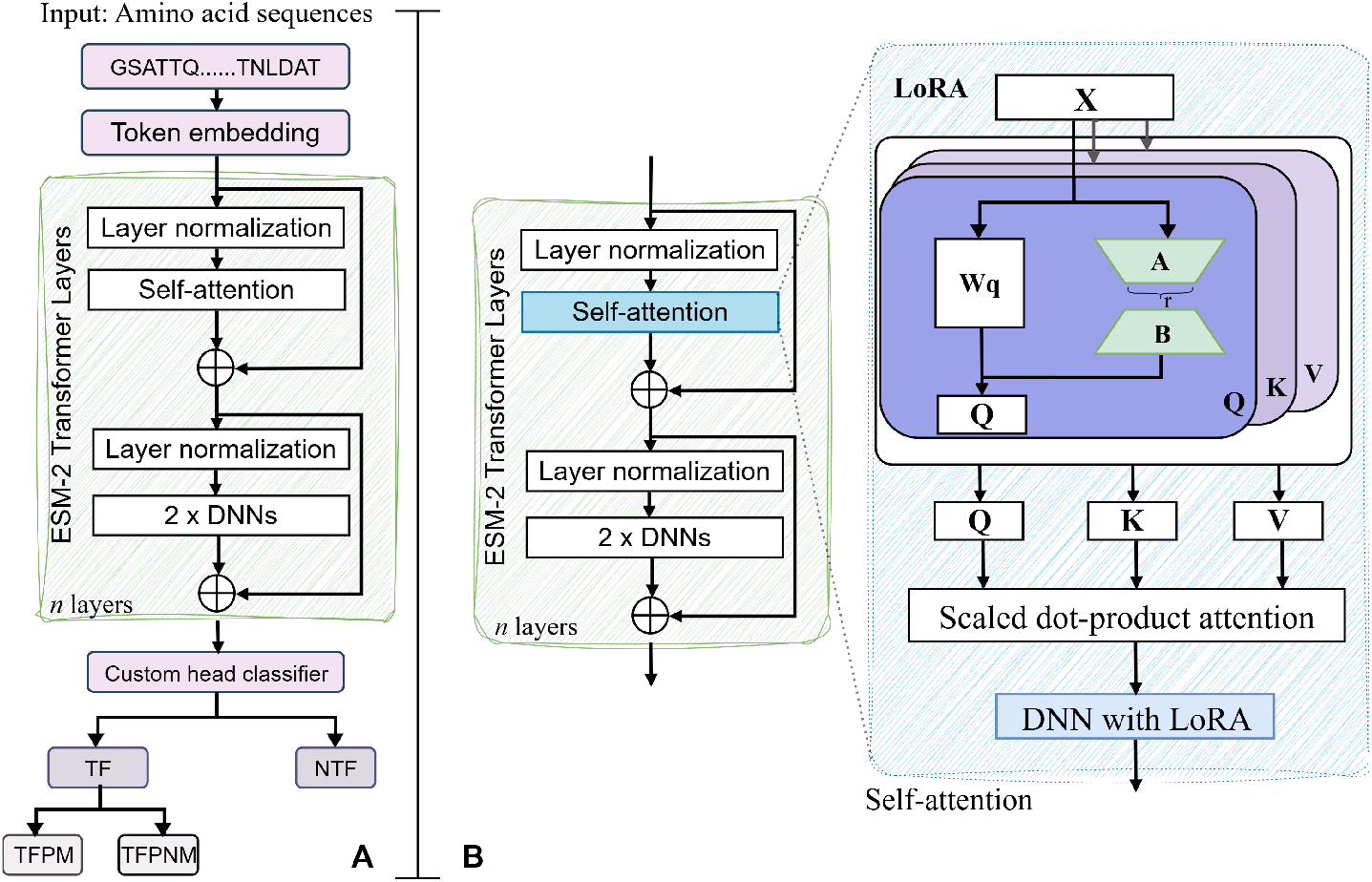
The architecture of the full and PEFT fine-tuning in the ESM2 models. (A) The ESM-2 backbone model uses the protein sequences to identify TF and NTF initially, and subsequently separates TF into TFPM and TFPNM categories. The *n* layers represent of the number of layers/depth of the model, where *n* = 6, 12, 30, 33, or 36 depending on the architecture of the ESM-2 model. (B) In the PEFT framework, LoRA introduces additional trainable low-rank matrices (r) into the self-attention module, enabling efficient fine-tuning without modifying the full model parameters.

To further interpret model behavior, we compare the attention weights from different layers and heads with known top motifs. This analysis aims to reveal how the model’s attention aligns with biologically relevant sequence patterns, providing insight into which residues and motifs are prioritized during transcription factor classification and interaction predictions. This enhanced understanding links computational attention patterns to established biological motifs, shedding light on the mechanistic basis of model decisions.

## Methods and material

### Data

The benchmark training and testing datasets were sourced from the study by Liu et al. [21]. The original dataset contained 601 human and 129 mouse transcription factors (TFs) with affinity for methylated DNA [27, 6], along with 286 TFs preferring non-methylated DNA [3]. To improve dataset quality, Liu et al. (TF dataset 2020) [21] applied several preprocessing steps. Sequences containing ambiguous amino acids such as “X”, “B”, or “Z” were first removed, followed by redundancy reduction using the CD-HIT clustering tool [28, 29] at a 25% sequence identity threshold. Additionally, sequences shorter than 50 residues were excluded. After these refinements, 522 TFs were retained as positive samples. For negative samples, an equal number of non-TFs were randomly selected from the UniProt database under the following criteria: reviewed status, experimental evidence at the protein level, sequence length greater than 50 amino acids, and absence of DNA-binding transcriptional activity. These selected sequences also exhibited less than 25% identity, as ensured by CD-HIT. Thus, the final TF dataset comprised 522 positive and 522 negative entries.

For constructing the first-layer model, identifying transcription factors against non-transcription factors (TF vs NTF), the TF dataset was divided into training and independent test sets at an 80:20 ratio. The TF training set consisted of 416 positive and 416 negative sequences, while the TF independent set contained 106 positives and 106 negatives.

For the second-layer model, targeting the identification of TFs with a preference for methylated DNA (TFPM dataset), the TFPM training set included 146 non-methylation-associated TFs (negative class) and 146 methylation-associated TFs (positive class). The TFPM independent test set consisted of 37 negatives and 69 positives.

### Feature Encoding

Feature extraction, or feature encoding, is a critical step in building robust machine learning models and directly influences their predictive performance. Among various approaches, evolutionary information captured through position-specific scoring matrix (PSSM) [30] profiles provides a richer representation than raw sequence data alone. Consequently, PSSM-based feature descriptors have become widely adopted as essential primary inputs for model construction, bridging an important gap in bioinformatics research. These descriptors have consistently enhanced prediction accuracy in diverse protein-related tasks, including fold recognition, structural class prediction, protein-protein interaction analysis, subcellular localization, RNA-binding site identification, and functional annotation. We extracted PSSM matrices using PSI-BLAST [31] run against UniProt-50 for each protein sequence. Each protein sequence of length *L* yielded a matrix of dimensions *L ×* 20. We processed the extracted PSSM matrices using row and/or column transformations and converted them in a 1D feature vectors representing different properties of the original PSSM matrix. In total we extracted seven different types of PSSM profiles and are defined as:

a. AAC PSSM: This feature, known as the auto-covariance transformation, is computed for column *j* by first calculating the mean of the column. Next, the deviation from this mean is obtained for the elements at rows *i* and *i* + *g*, which are then multiplied together. The summation is performed by iterating *i* from 1 to *L* − *g*. Since *j* ranges from 1 to 20 and *g* ranges from 1 to 10, the resulting representation yields a feature vector of length 200, as defined below:

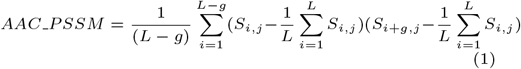

where *L* is the length of the protein sequence, *S*_*i*,*j*_ represents the PSSM score at position *i* for amino acid *j*. This feature captures the dependency and spatial correlation of evolutionary information along the sequence, enhancing the representation of protein characteristics.
b. AB_PSSM: This feature encompasses two distinct types of feature vectors. Initially, each protein sequence is segmented into 20 equally sized sections, referred to as blocks, and within each block, the corresponding row vectors from the PSSM are summed. The resultant vector for each block is then normalized by dividing by the block’s length, which corresponds to 5% of the total protein length. Arranging these 20 normalized vectors sequentially yields the first feature vector with a length of 400. For the second type, the mean of the positive values in each column of the PSSM is calculated for each block and each amino acid. These 20 average values, one for each block, are concatenated for each of the 20 amino acid types, producing a vector of length 20 per amino acid; combining these across all amino acids results in a second feature vector of length 400.
c. Trigram PSSM: This feature vector, extracted from the PSSM, has a length of 8,000. It is computed by multiplying elements from three consecutive rows and three distinct columns of the PSSM. This operation is applied across all possible sets of three consecutive rows, and the resulting products are summed to yield a single feature corresponding to the selected three columns. Since there are 20 possible columns, the total length of the feature vector becomes 20 *×* 20 *×* 20 = 8,000. It can be defined as:

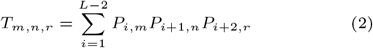

where *P*_*i,m*_ represents the PSSM score at row *i* and column *m, L* is the length of the protein sequence, and *m, n, r* ∈ *{*1, …, 20*}* correspond to the amino acid columns.
d. DP PSSM: This feature, associated with dipeptide composition (DPC), was initially introduced for predicting the structural class of proteins. To compute this descriptor, elements from two consecutive rows and two distinct columns in the PSSM are multiplied together. This multiplication is performed across all relevant pairs of rows and columns. It can be defined as:

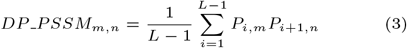

where *P*_*i,m*_ and *P*_*i*+1,*n*_ are elements from two consecutive rows (i and i+1) and two distinct columns (m and n) of the PSSM matrix, and *L* is the length of the protein sequence.
e. PSe PSSM: This feature was initially introduced for the prediction of membrane proteins and their respective types [32]. The PSe PSSM descriptor forms a 320-dimensional vector, where the first 20 elements represent the average values of the 20 rows in the PSSM matrix. The remaining components are calculated as follows: for each column, the mean squared difference between the *i*^*th*^ and (*i* + *lag*)^*th*^ elements is computed, with *lag* taking integer values from 1 to 15. Consequently, the total length of the resulting feature vector is 20 *×* 15 + 20 = 320 The process of generating this feature is expressed in Equation 4.

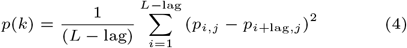

where *j* = 1, 2, …, 20, lag = 1, 2, …, 15*k* = 20 + *j* + 20(lag − 1)
f. Smoothed PSSM: Smoothed PSSM refers to a representation of a position-specific scoring matrix in which local sequence context is incorporated by averaging or summing evolutionary scores over a window of neighboring residues. Instead of using the raw scores for each position independently, a smoothing (or sliding) window is applied to the PSSM so that each position’s score reflects not only its own evolutionary information but also that of its surrounding residues. This approach captures local patterns and contextual conservation, which can improve prediction of functional regions or protein attributes. If the window extends beyond sequence boundaries, zero vectors are typically appended to maintain consistency in the profile length. For a residue at position *i*, the Smoothed PSSM value is calculated by incorporating the scores of neighboring residues within a specified window. Mathematically, this can be represented as:

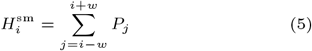

where *w* denotes the window radius and *P*_*j*_ is the PSSM score at position *j*. If the summation window extends beyond the sequence termini, zero-padding is applied to maintain the length of the resulting profile vector.
g. *K* separated bigram PSSM: *K* separated bigram PSSM is a feature extracted from the Position Specific Scoring Matrix, which extends the concept of dipeptide composition (DPC). For each pair of distinct columns *m* and *n*, it considers pairs of residues separated by a fixed gap of length *k* along the sequence. Mathematically, the feature for columns *m* and *n* with separation *k* is calculated by:

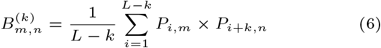

where *P*_*i,m*_ and *P*_*i*+*k,n*_ are PSSM scores at positions *i* and *i* + *k* for columns *m* and *n*, respectively, and *L* is the protein length. The special case when *k* = 1 corresponds exactly to the dipeptide composition (DPC) feature.

Apart from the PSSM profiles, we extracted sequence-level embeddings from the pre-trained models and used the same as an input to a 128 node custom-head classifier. The embeddings were extracted from the last hidden layer of a model.

### Pre-trained pLMs

The pLMs employed in this study varied considerably in both architecture and scale (Table 1), ranging from 8 million parameters (ESM2-8M) to 3 billion parameters (ESM2-3B). The ESM2 models are encoder-only architectures based on RoBERTa64 and trained with an unsupervised masked language modeling objective [33]. In contrast, the other pLM, Prot-Bert, is built on BERT architecture [34], an encoder large language model pretrained using a span-masking objective [35, 34]. All models were initialized from pre-trained checkpoints publicly available on HuggingFace. The pre-trained models were utilized to extract the sequence-level embeddings from the last hidden layer. The size of the sequence-level embeddings is mentioned in Table 1. Extraction of embeddings from the pre-trained ESM2-3B model failed due to the model exceeding available memory limits, even with an effective batch size of 1. The final classification task was carried out by a 128 node two layer custom head classifier and the same setting was carried on the all pre-trained models for the standard evaluation.

**Table 1.**
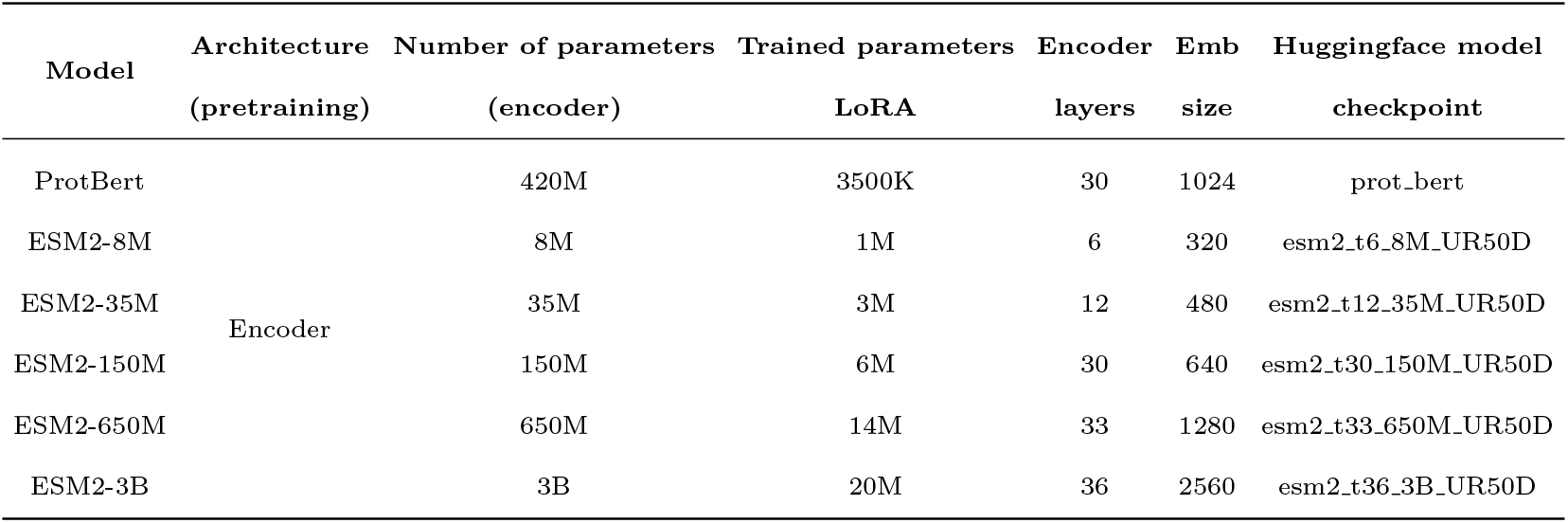
Details of the models and their respective embedding size.

### Model training

Top-level comparisons were made between pre-trained and fine-tuned models as follows. For the pre-trained results, we generated embeddings for all sequences. For tasks operating at the protein level, sequence embeddings were reduced to a single fixed-length vector by averaging across the sequence dimension, preserving the original embedding size. This vector was then passed through a prediction head, a custom head classifier with a fully connected layer containing 128 neurons. Training stopped when the plateau in training loss was reached, with each experiment repeated five times using different random seeds.

For fine-tuning, we attached an identical fully connected layer (size 128) to each pLM encoder to serve as the prediction head. In the case of ProtBert, the final hidden states were average-pooled across the sequence length during training for protein-level tasks. Following the recommendations of the ESM2 authors [36], the prediction head for ESM2 was linked exclusively to the first token (the special <CLS> token). Each experiment was run three times with different random seeds, and training was stopped once both the training and validation losses plateaued. To improve computational efficiency, Parameter-Efficient Fine-Tuning (PEFT) was applied across all models using the Low-Rank Adaptation (LoRA) approach [37]. For the smaller ESM2 variants (up to the 650M model), we additionally evaluated full fine-tuning of the entire network

For both embedding-based and fine-tuning approaches, we reported the performance of the checkpoint yielding the lowest validation loss for each run. The improvements were calculated as:

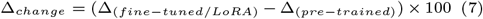

where Δ_(*k*)_ represents the best performance of the model *k*.

All models and embedding extraction were carried out on an Nvidia GeForce RTX 3080 GPU coupled with an Intel i9 processor with 128GB memory using PyTorch 2.2.0+cu118. The GPU is good enough to run and train all models but full fine-tuning ESM2 3B due to its sheer size.

### Evaluation metrics

Machine learning, deep learning, and graph neural networks find widespread application in bioinformatics [38, 39, 40, 41, 42], chemoinformatics [43, 44], and image analysis [45, 46, 47]. Evaluating the performance of computational models is critical to establish their predictive accuracy and reliability. To this end, a standard suite of evaluation metrics is employed, including accuracy (Acc), specificity (Sp), sensitivity (Sn), precision (Pr), F1 score (F1), area under the receiver operating characteristic curve (AUC), and Matthews correlation coefficient (MCC). These metrics are extensively utilized in bioinformatics research for providing a comprehensive understanding of model behavior, highlighting both strengths and weaknesses [48, 49, 50, 51, 52]. Definitions of these metrics are provided as follows:

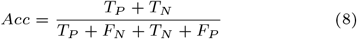

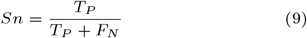

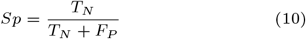

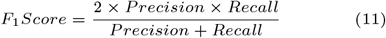

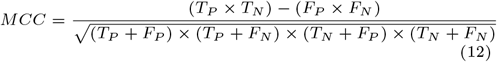

where *T*_*P*_ represents true positive, *T*_*N*_ represents true negative, *F*_*P*_ represents false positive, and *F*_*N*_ represents false negative.

## Results

### Performance of baseline models on raw PSSM features

We assessed the performance of each PSSM-based feature representation individually across five distinct machine learning classifiers—RF, XGB, ETC, LGBM, and CatBoost—using a 5-fold cross-validation strategy. Seven feature encodings—AAC PSSM, AB PSSM, Tri gram PSSM, DP PSSM, Smoothed PSSM, *K* separated bigram PSSM, and PSe PSSM—were evaluated against these five conventional machine learning classifiers, frequently used in the field of bioinformatics [53, 54, 55], in the identification of transcription factors, and the detailed results are mentioned in Supplementary Tables S1 and S2. From the Figure 2, the cross-validation MCC is in the range of 0.307-0.805, showing variable results across different feature encodings. Among the seven PSSM features, Trigram PSSM shows the highest CV-MCC value of 0.805 against the ETC classifier and the least CV-MCC score of 0.602 against the XGB classifier, thus showing the variation of almost 20% in the CV-MCC scores across different classifiers. The DP PSSM shows consistent performance with a little variation across the different classifiers, with values ranging from 0.726-0.750. The Smoothed PSSM encoding was the least effective feature encoding type irrespective of the type of the classifier, with CV-MCC ranging from 0.307-0.580. Among the different classifier algorithms, the XGB model was the least performing one, with values ranging from 0.307-0.743 and the ETC showed better results than the other classifiers against all PSSM features but two.

**Figure 2.**
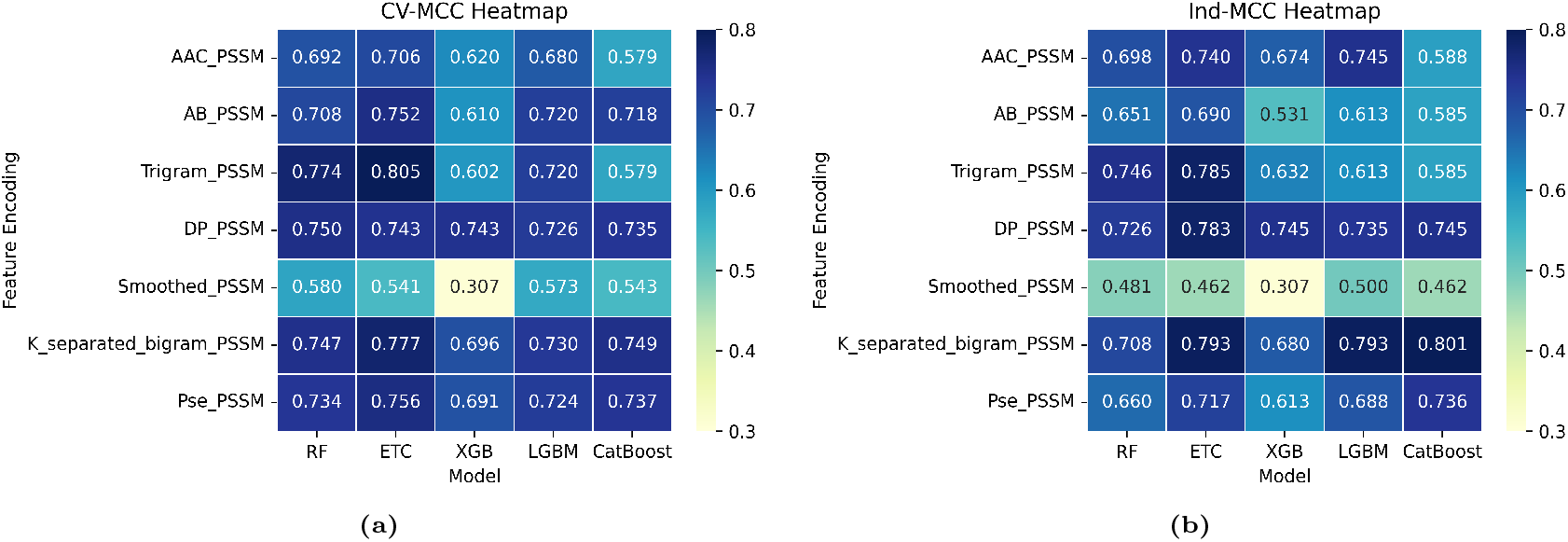
Heatmap representing MCC score of seven PSSM feature encodings against five machine learning models (a) cross-validation MCC score (b) independent MCC score

On the independent dataset, the classifiers show the similar pattern as can be seen from the Figure 2. The Smoothed PSSM showed the least performance across the different classifiers, with values ranging from 0.307-0.500, akin to the cross-validation performance. Here again, DP PSSM shows the consistent performance with values ranging from 0.726-0.783. The CatBoost classifier showed the highest independent MCC value of 0.801 against the *K* separated bigram PSSM. The XGB model showed the least performance among the five classifiers, with values ranging from 0.307-0.745. From Supplementary Tables S1 and S2, all PSSM features except Smoothed PSSM showed consistent performance in terms of accuracy across the different classifiers on the main benchmark and independent datasets.

### Performance of the pre-trained models

Before fine-tuning the protein-language models, we extracted the embeddings from five models of moderate size, which include ESM2-8M, ESM2-35M, ESM2-150M, ESM2-650M, and ProtBert. For evaluation, we constructed a custom classifier head of a 128-D dense layer, rather than using the default classifier head provided by the Hugging Face, for more control and consistency. The performance of the custom head classifier on the independent dataset embeddings showed that the ESM2-650M model is performing best as compared to the other models, which is understandably due to the 650M parameter size, the highest among these five protein models (Table 2). The custom head classifier achieved an accuracy of 0.932±0.003 and an MCC of 0.831±0.038, and the ESM2-8M model showed the least performance as compared to the other models, achieving the best accuracy of 0.857±0.008 and an MCC value of 0.716±0.014, reflecting its meager parameter size as the main reason for its below-par performance. The ProtBert model was somewhere intermediate in performance as compared to the other four models with an accuracy of 0.906±0.220 and an MCC score of 0.814±0.039.

**Table 2.** Performance comparison of the pre-trained models on the independent TF set.

| Model | Parameters | Acc | Sn | Sp | MCC | AUC | F1 |
| --- | --- | --- | --- | --- | --- | --- | --- |
| ESM2 8M | 8M | $0.857 \pm 0.008$ | $0.890 \pm 0.017$ | $0.824 \pm 0.026$ | $0.716 \pm 0.014$ | $0.936 \pm 0.016$ | $0.861 \pm 0.006$ |
| ESM2-35M | 35M | $0.890 \pm 0.004$ | $0.913 \pm 0.017$ | $0.868 \pm 0.026$ | $0.782 \pm 0.008$ | $0.958 \pm 0.001$ | $0.867 \pm 0.025$ |
| ESM2-150M | 150M | $0.924 \pm 0.000$ | $0.934 \pm 0.001$ | $0.915 \pm 0.028$ | $0.840 \pm 0.038$ | $0.970 \pm 0.011$ | $0.920 \pm 0.020$ |
| ESM2-650M | 650M | $0.932 \pm 0.003$ | $0.928 \pm 0.010$ | $0.937 \pm 0.005$ | $0.831 \pm 0.038$ | $0.972 \pm 0.006$ | $0.916 \pm 0.020$ |
| ProtBert | 450M | $0.906 \pm 0.020$ | $0.934 \pm 0.024$ | $0.878 \pm 0.032$ | $0.814 \pm 0.039$ | $0.969 \pm 0.012$ | $0.908 \pm 0.020$ |

### Performance of the full fine-tuned models

All five models were fully fine-tuned along with the custom head classifier of the 128-D dense layer to enhance its prediction and interpretation. Each model ran for a total of 10 epochs, and the effective batch size was set as 8. For larger models we used the gradient accumulation steps parameter to avoid out-of-memory issues due to resource constraints. The training accuracy-loss curve is shown in Figure 3. It can be seen from Figure 3, the training loss and the validation loss do converge in all cases of the ESM2-based models. The performance of the five protein-language models on the independent dataset is mentioned in Table 3. The ESM2-650M model achieved an accuracy of 0.948±0.015 and an MCC score of 0.898±0.030, the highest among the five pLMs. The ESM2-8M model, with the least number of parameters, also showed good performance, achieving an accuracy of 0.910±0.019 and an MCC value of 0.820±0.038. To assess the stability of the ESM2 models, we employed bootstrapping with 1,000 resampled test sets and evaluated performance across each iteration. The bootstrapped performance results demonstrate that the ESM2 models are both robust and highly stable across repeated resampling of the test set. The narrow standard deviations observed for all key metrics indicate that the model’s behavior does not fluctuate significantly with changes in sample composition. This low variability confirms that the model’s predictions are not driven by chance patterns or specific subsets of the data, but instead reflect consistently reliable discrimination between transcription factor and non-transcription-factor sequences.

**Table 3.** Performance comparison of the fully fine-tuned models on the independent TF dataset.

| Model | Parameters | Acc | Sn | Sp | MCC | AUC | F1 |
| --- | --- | --- | --- | --- | --- | --- | --- |
| ESM2-8M | 8M | $0.910 \pm 0.019$ | $0.923 \pm 0.026$ | $0.896 \pm 0.028$ | $0.820 \pm 0.0381$ | $0.945 \pm 0.016$ | $0.911 \pm 0.019$ |
| ESM2-35M | 35M | $0.924 \pm 0.018$ | $0.923 \pm 0.026$ | $0.925 \pm 0.026$ | $0.848 \pm 0.036$ | $0.959 \pm 0.014$ | $0.924 \pm 0.019$ |
| ESM2-150M | 150M | $0.939 \pm 0.016$ | $0.971 \pm 0.016$ | $0.906 \pm 0.028$ | $0.879 \pm 0.031$ | $0.974 \pm 0.012$ | $0.940 \pm 0.016$ |
| ESM2-650M | 650M | $0.948 \pm 0.015$ | $0.972 \pm 0.016$ | $0.925 \pm 0.026$ | $0.898 \pm 0.030$ | $0.979 \pm 0.009$ | $0.949 \pm 0.015$ |
| ProtBert | 450M | $0.501 \pm 0.033$ | $1.000 \pm 0.000$ | $0.000 \pm 0.000$ | $0.000 \pm 0.000$ | $0.500 \pm 0.000$ | $0.666 \pm 0.029$ |

**Figure 3.**
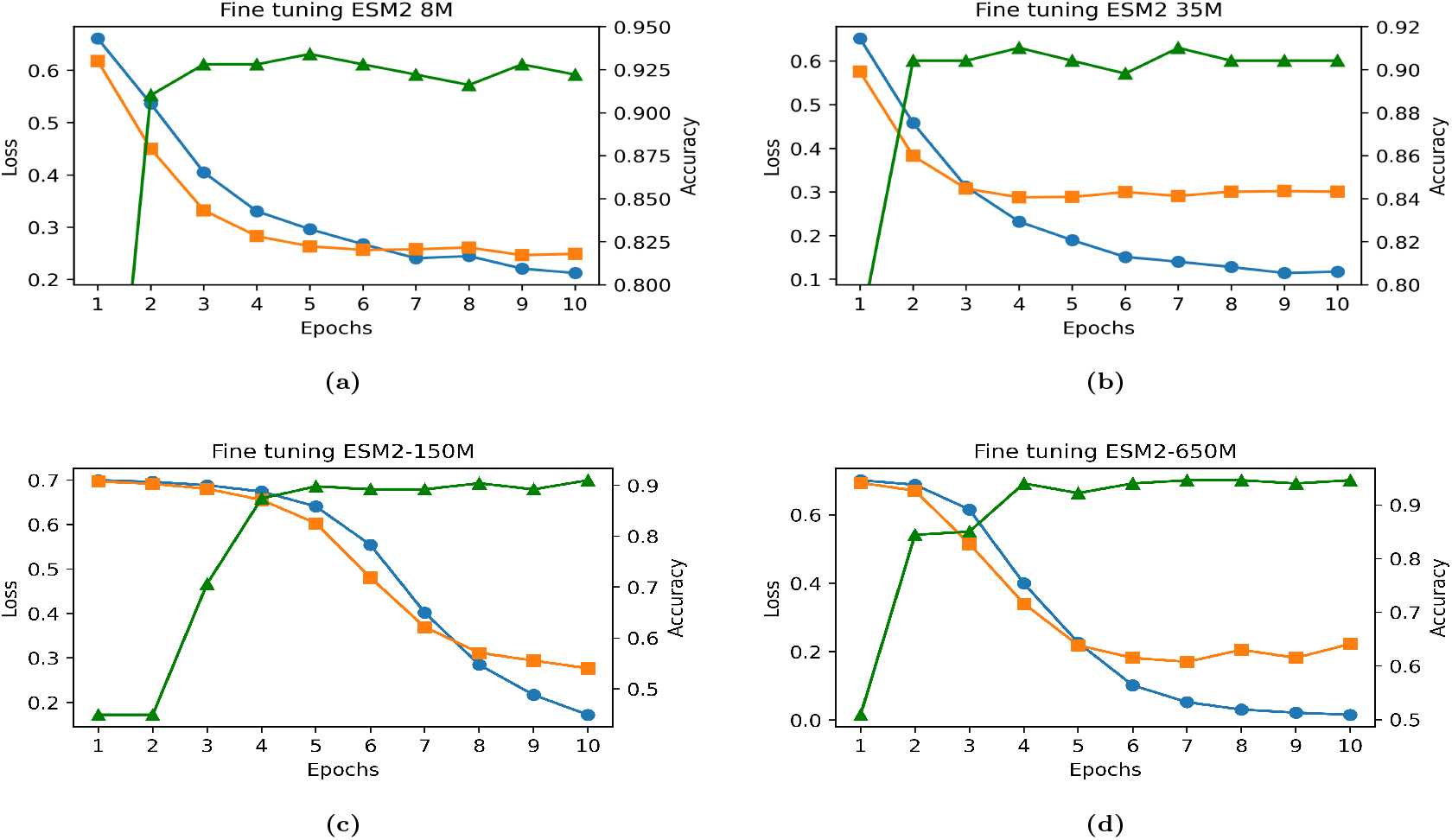
Training accuracy-loss curve of the four evolutionary pLMs. The green represents training accuracy, blue represents the training loss, and orange represents the validation loss curve. (a) accuracy-loss curve of the full fine-tuned ESM2-8M (b) accuracy-loss curve of the full fine-tuned ESM2-35M (c) accuracy-loss curve of the full fine-tuned ESM2-150M (d) accuracy-loss curve of the full fine-tuned ESM2-650M

The performance of the ProtBert, a 450M model, is quite disappointing, as the model was not able to learn anything even with hyperparameter tuning, with the training loss curve remaining flat, thus showing BERT-based models’ inability to learn any meaningful information or features. The results demonstrated (Table 3) that the fine-tuned models, except ProtBert, showed significant improvements over the pre-trained models (Table 2). The improvement in the fine-tuned models is largely because of the inherent nature of the large language models to learn the intrinsic characteristics of the protein sequences.

### Using LoRA

Apart from the fine-tuning full layers, we used the PEFT technique to understand the impact on the performance of the protein-language models. We tried LoRA with variable rank values ranging from 4 to 32. The effective batch size was again kept at 8, the same as in fully fine-tuned models, for effective comparison. The LoRA technique does reduce the training time and GPU memory usage considerably but has limited impact on the performance of the protein-language models. The performance of the protein-language models using LoRA is depicted in Table 4. The LoRA doesn’t bring any significant improvement in the performance of the protein models over fully fine-tuned models, as can be seen from Tables 3 and 4. Again, the ProtBert models don’t show any learning process even with LoRA and hyperparameter optimization, thus revealing their inability to learn anything because of their BERT-based structure. The Evolutionary Scale Modeling 2-based models showed a good learning process, with the ESM2-650M achieving the highest accuracy of 0.943±0.016 and the highest MCC score of 0.887±0.032. The ESM2-8M was the least performing model with an accuracy of 0.900±0.020 and an MCC score of 0.801±0.041, slightly less than the fully fine-tuned corresponding model. With LoRA using a fraction of the original parameters, we were able to train the ESM2-3B model as well, achieving the highest accuracy of 0.934±0.017 and an MCC score of 0.868±0.034, with the rank set as 4, thus showing the ability of the LoRA technique to run large language models on small GPUs.

**Table 4.** Performance assessment of LoRA models on the independent TF dataset.

| Model | Parameters | Acc | Sn | Sp | MCC | AUC | F1 |
| --- | --- | --- | --- | --- | --- | --- | --- |
| ESM2-8M | 8M | 0.900 ± 0.020 | 0.895 ± 0.030 | 0.906 ± 0.028 | 0.801 ± 0.041 | 0.935 ± 0.018 | 0.900 ± 0.022 |
| ESM2-35M | 35M | 0.924 ± 0.018 | 0.923 ± 0.026 | 0.925 ± 0.026 | 0.849 ± 0.036 | 0.951 ± 0.017 | 0.924 ± 0.019 |
| ESM2-150M | 150M | 0.939 ± 0.016 | 0.943 ± 0.022 | 0.934 ± 0.024 | 0.878 ± 0.033 | 0.961 ± 0.015 | 0.939 ± 0.017 |
| ESM2-650M | 650M | 0.943 ± 0.016 | 0.934 ± 0.024 | 0.953 ± 0.021 | 0.887 ± 0.032 | 0.982 ± 0.008 | 0.943 ± 0.016 |
| ESM2-3B | 3B | 0.934 ± 0.017 | 0.924 ± 0.026 | 0.944 ± 0.022 | 0.868 ± 0.034 | 0.981 ± 0.009 | 0.933 ± 0.017 |
| ProtBert | 450M | 0.501 ± 0.033 | 1.00 ± 0.000 | 0.000 ± 0.000 | 0.000 ± 0.000 | 0.500 ± 0.000 | 0.667 ± 0.029 |

The improvements in the fully fine-tuned and LoRA models over the pre-trained models are depicted in Figure 4. The improvements were calculated as per the equation 7. The fully fine-tuned and LoRA models showed improvements over the pre-trained models across different metrics (Figure 4). The fully fine-tuned models showed overall improvement across all metrics over pre-trained models. The ESM2-8M and ESM2-35M models were able to show variable improvements across different metrics over pre-trained models. The ESM2-150M and ESM2-650M models also showed modest improvements across all metrics but specificity. The specificity values decreased by 0.94% in the ESM2-150M model and 1.25% in the ESM2-650M model. In the case of LoRA models, the largest overall gain occurs in the ESM2 8M model, especially in specificity (+8.15%) and MCC score (+8.47%). The ESM2-35M shows strong balanced gains, notably in F1 (+5.70%) and MCC (+6.65%). The ESM2-150M and ESM2-650M also showed improvements, but gains are smaller, with modest improvements spread across all metrics. The AUC improves only for ESM2-650M (+0.61%), while it slightly decreases in ESM2-35M and ESM2-150M. The low standard deviations indicate that the performance of ESM2 models are stable across different resampled versions of the test set, meaning the results are not overly sensitive to which samples happen to be in the test split. The accuracy-loss curves of the LoRA ESM2 models are depicted in Figure 5.

**Figure 4.**
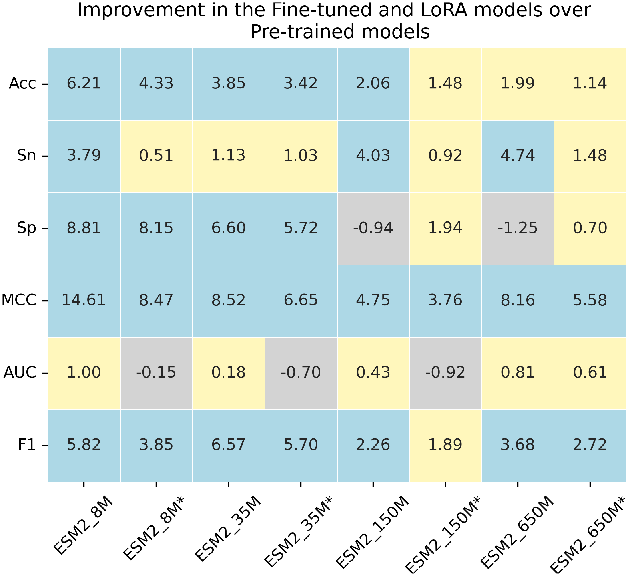
The values represent improvement in the fine-tuned and LoRA models over Pre-trained models in percentage terms. The blue tiles represent the positive improvement of above 2%, yellow tiles represent improvement of 0%-2%, and purple tiles represents the negative improvement. * denotes the LoRA models.

**Figure 5.**
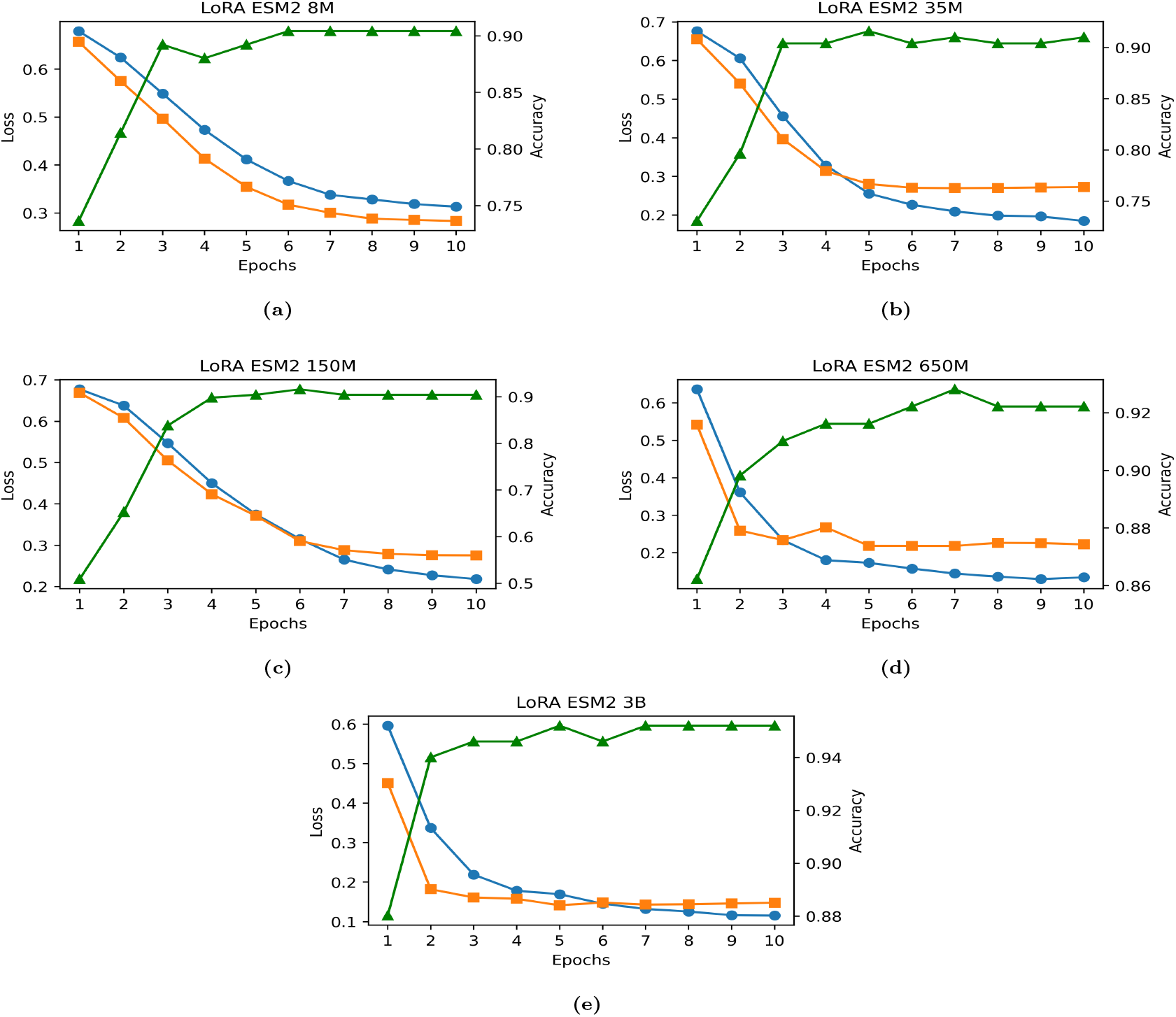
Training accuracy-loss curve of the five ESM2 models using PEFT technique LoRA. The green represents training accuracy, blue represents the training loss, and orange represents the validation loss curve. (a) accuracy-loss curve of the LoRA ESM2-8M (b) accuracy-loss curve of the LoRA ESM2-35M (c) accuracy-loss curve of the LoRA ESM2-150M (d) accuracy-loss curve of the LoRA ESM2-650M (e) accuracy-loss curve of the LoRA ESM2-3B

The primary factors influencing computational resource demands during model training were the parameter sizes of the pLMs and the quadratic scaling of attention mechanisms, which increased requirements for longer protein sequences. Modern GPUs with over 10GB of memory are generally capable of handling all pLMs evaluated in this study, except the ESM2-3B model. With these approaches, most pLMs could be fine-tuned even on older GPUs with as little as 10GB of memory. Interestingly, both full fine-tuning and parameter-efficient LoRA fine-tuning required similar amounts of GPU memory for smaller models, differing only in training speed (Figure 6). In contrast, generating embeddings required much less GPU memory, making it suitable for datasets with very long sequences. The runtime values of the fully fine-tuned and LoRA models are mentioned in Supplementary Table S3.

**Figure 6.**
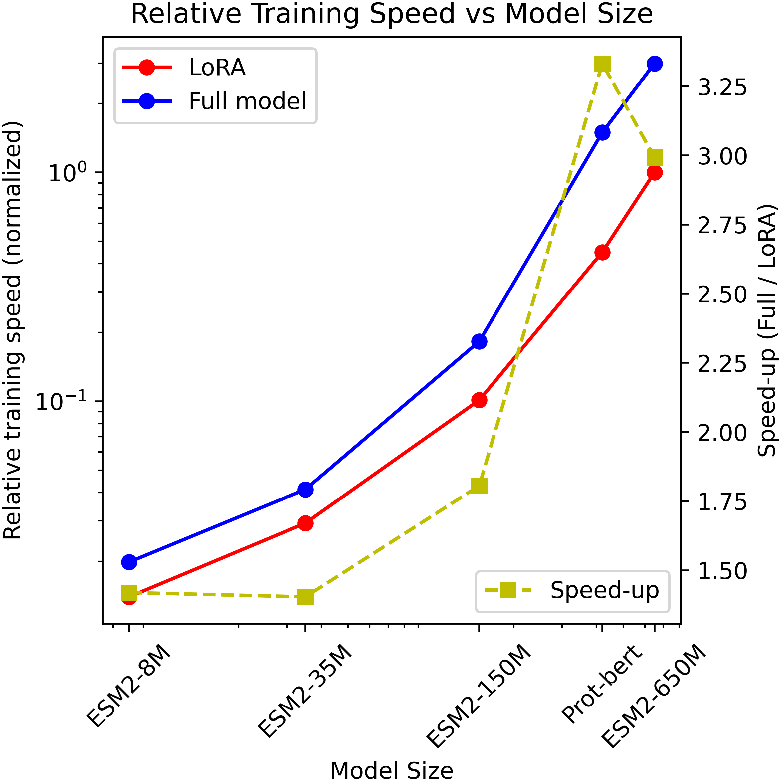
Fine-tuning training speed. The training speeds of full fine-tuning (blue) and LoRA (red) are displayed on a normalized scale, using ESM2-650M LoRA fine-tuning as the reference point with a baseline value of 1 (x). The corresponding speed-up achieved by each model (olive) is shown on a linear scale. All experiments used transcription factor protein sequences of length 1024 in a per-protein training setup. The LoRA technique is more beneficial as it runs faster than the full fine-tuned models. In the case of the largest model, ESM2-650M, LoRA fine-tuning was approximately 3 times faster than full fine-tuning.

### Performance evaluation with the existing predictors

We evaluated the performance of our fully fine-tuned ESM2-650M model with the existing state-of-the-art predictors in the literature, namely TFPred [21], Li RNN [25], PSSM+CNN [22], Capsule TF [23], and TFProtBert [26]. From Table 5, it is clear that the full fine-tuned model, ESM2-650M, is achieving the best possible results among the existing methods, with an accuracy of 0.948, which is higher than the rest of the lot in the range of 3.6%-11.8%, and with an MCC score of 0.899, which is also significantly higher than the existing methods, thus showing the robust nature of the full fine-tuned model. The fully fine-tuned model also has the highest sensitivity, specificity, and ROC-AUC values among the other models (Table 5).

**Table 5.** Performance comparison between the existing models and the best fully fine-tuned model on the independent dataset. ^*†*^represents the full fine-tuned model.

| Method | ACC | Sn | Sp | MCC | AUC |
| --- | --- | --- | --- | --- | --- |
| TFPred | 0.830 | 0.802 | 0.859 | 0.661 | 0.912 |
| Li-RNN | 0.867 | 0.887 | 0.840 | 0.727 | 0.913 |
| PSSM+CNN | 0.873 | 0.906 | 0.840 | 0.746 | 0.960 |
| Capsule_TF | 0.882 | 0.915 | 0.850 | 0.765 | 0.925 |
| TFProtBert | 0.912 | 0.931 | 0.911 | 0.867 | 0.942 |
| ESM2-650M<sup>†</sup> | 0.948 | 0.972 | 0.925 | 0.899 | 0.979 |

### Second layer performance

In the second layer, the transcription factors as predicted in the first layer were further classified into transcription factors preferring binding to the methylated DNA (TFPM) and the transcription factors preferring binding to the non-methylated DNA (TFPNM). The second layer performance of the full fine-tuned and the LoRA models using ESM2 variants on the independent dataset (TFPM independent set) is mentioned in Supplementary Tables S4 and S5, respectively. The performance of the ESM2-8M full fine-tuned model is slightly better than the LoRA 8M model, as can be seen from the Supplementary Tables S4 and S5. The ESM2-8M fully fine-tuned achieved the highest accuracy of 0.593 ± 0.043 and an MCC score of 0.149 ± 0.067, and the LoRA 35M model achieved the highest accuracy of 0.495 ± 0.046 and an MCC score of 0.111 ± 0.089.

The performance, on the TFPM independent dataset, of the fully fine-tuned ESM2-8M model against the existing methods, Li RNN [25] and TFProtBert [26], is shown in the Table 6. The fully fine-tuned ESM2-8M shows around 4%-29% improvement in the accuracy metric compared to the existing methods, whereas in the case of the MCC metric, the fully fine-tuned method shows improvement in the range of 8%-63%. These results show the robustness of the fully fine-tuned ESM2-8M model in identifying TFPMs and TFPNMs as compared to the existing methods, with a great scope of improvement.

**Table 6.** The performance comparison between the existing methods and the best model on the second layer in identifying TFPMs and TFPNMs. ^*†*^represents the fully fine-tuned model.

| Method | Acc | Sn | Sp | MCC | AUC |
| --- | --- | --- | --- | --- | --- |
| Li_RNN | 0.264 | 0.304 | 0.189 | -0.483 | — |
| TFProtBert | 0.556 | 0.594 | 0.486 | 0.077 | 0.537 |
| ESM2-8M<sup>†</sup> | 0.593 | 0.414 | 0.726 | 0.149 | 0.511 |

### Attention Maps and Motifs

To investigate how the top model’s attention mechanism captures functionally important amino acid fragments, we adapted a technique from prior work [56, 57]. We examined how the attention-based amino acid confidence scores from the best full fine-tuned model, ESM2-650M, aligned with the top known motifs extracted using the STREME web server [58] from the TF dataset (positive TF samples). As shown in Figure 7, the amino acids or fragments that received the highest attention corresponded closely to those that frequently occur in the motif. The top motif in the positive benchmark TF dataset identified by the STREME [58] is “ERKRRRNR”, and its corresponding attention map is depicted in Figure 7a. This suggests that the model’s attention mechanism effectively identifies biologically meaningful regions, providing a plausible explanation for its predictions. Furthermore, full fine-tuning further refines the model’s ability to capture functional interactions among amino acids. This results in a more distinct and varied attention pattern across relevant amino acid pairs, enhancing the model’s performance in sequence classification.

**Figure 7.**
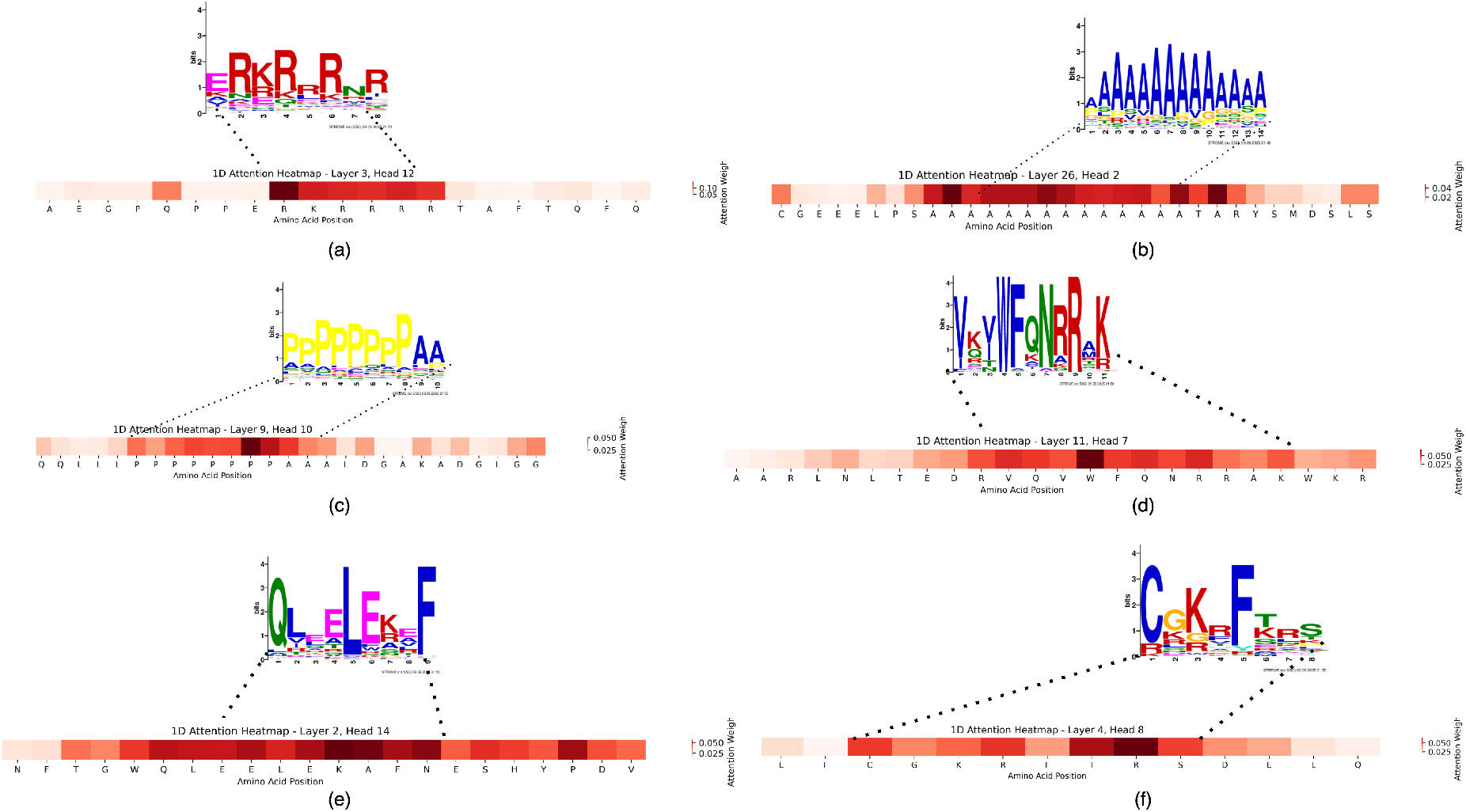
Visualization of attentions for various protein sequences corresponding to the top six motifs. The protein sequences are too long to be used for attention visualization, so we extracted only the high attention part from the original sequences corresponding to the motifs for better visualization. For each of the six highest-scoring motifs, the corresponding high-attention regions from representative protein sequences are extracted and shown with their 1D attention heatmaps. The top panels display the sequence logos generated from the attention-enriched subsequences, while the bottom panels show the amino-acid positions and their normalized attention weights across different transformer layers and heads.

Figure 8 depicts the attention pattern of the transcription factor protein sequence containing the top motif “ERKRRRNR”. The attention pattern, as shown in Figure 8a, depicts stronger attention (darker blue regions), occurring primarily in the middle of the sequence, especially among and around the repeated “RKRRRR” segment. This data indicates that these amino acids are being focused on more intensely by this attention head, either attending to themselves or to each other. Layer 8, Head 16 of the model appears to focus on and within the central, arginine/lysine-rich motif of the protein, likely attending to biologically significant features or repetitive sequence regions. This pattern suggests that this head plays a role in identifying related motifs, which are characterized by clusters of basic residues. Figure 8b shows the darkest blue regions are concentrated strictly along the main diagonal, indicating that most of the attention in this head is focused on each amino acid attending to itself or its immediate neighbors. Unlike Layer 8, Head 16, which captured strong inter-residue attention, especially around repetitive motifs, Layer 11, Head 3 in this map predominantly reinforces local or self-attention, acting conservatively rather than extracting or aggregating broader features. Together, the two heads reveal complementary attention behaviors, one specialized in motif-level enrichment and the other maintaining global positional structure.

**Figure 8.**
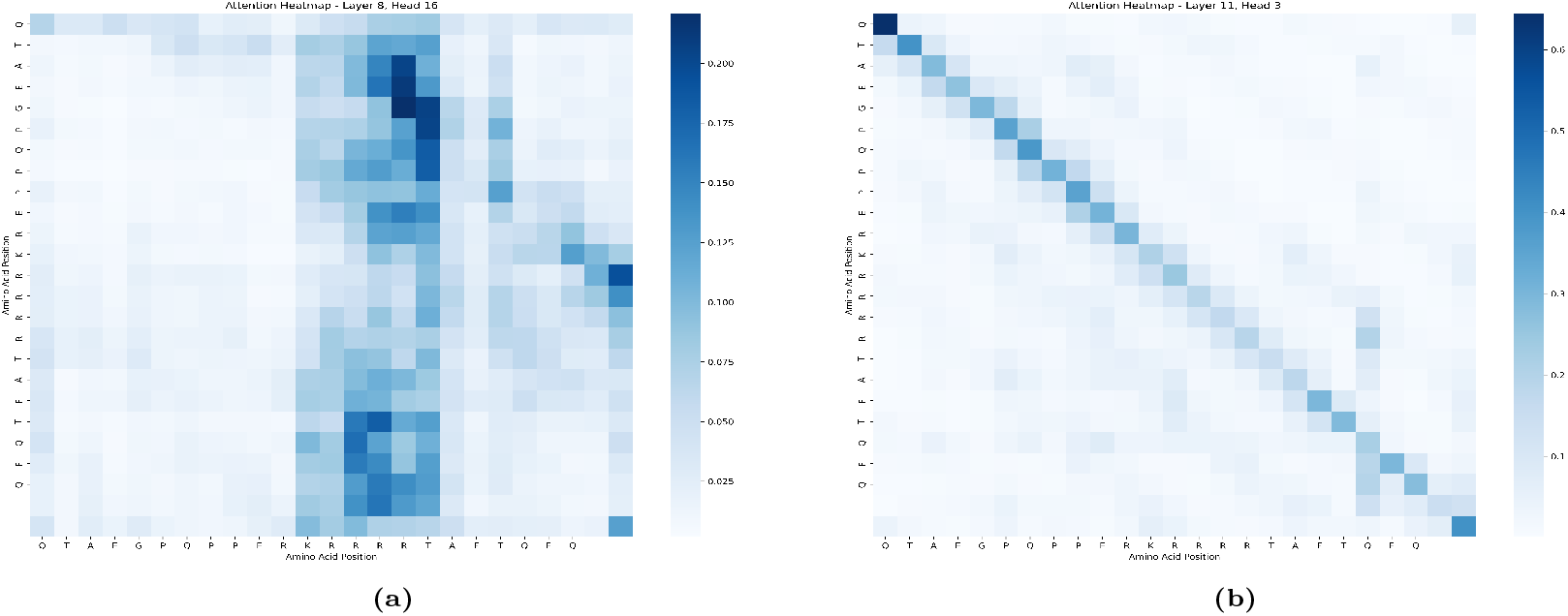
The attention heatmap corresponding to the protein sequence containing the top motif “ERKRRRNR” shows the nature of the attention pattern in different layers and heads. (a) Layer 8, Head 16 shows a strong localized attention concentration around the motif region, indicating that this head specifically captures the arginine-rich segment and its surrounding contextual residues (b) Layer 11, Head 3 displays a more diagonal, self-attention–dominant pattern, reflecting position-wise identity attention while still highlighting mild emphasis around the motif.

## Conclusion

In this research, we utilized the pLM ESM2 models (Evolutionary Scale Modeling 2) and ProtBert. The comparison between pre-trained models, full fine-tuned models, and LoRA models revealed the fact that the full fine-tuned and LoRA models achieved better results than the pre-trained models on the account of their ability to adapt more effectively to downstream tasks through fine-tuning. This adaptation allows them to specialize and improve performance on specific target tasks compared to the generalist pre-trained models, which are trained on broad data and not specifically optimized for those tasks. Fine-tuning, including LoRA’s parameter-efficient approach, enables updating model parameters to capture task-specific patterns, thus improving task accuracy and generalization beyond what is achievable by just using the pre-trained representations alone. The full fine-tuning is resource- and time-intensive, whereas LoRA is resource-effective, and the time complexity is slightly better than the full fine-tuning. The full fine-tuned ESM-650M model, on the TF independent set, achieved the highest accuracy of 0.948 ± 0.015 and an MCC score of 0.898 ± 0.030, while the LoRA ESM2-650M model reached an accuracy of 0.943 ± 0.016 and an MCC of 0.887 ± 0.032, showing only slight performance variation. Notably, the LoRA model completes training in less than one-third of the time required by full fine-tuning, compensating for its marginally lower accuracy. This advantage is more pronounced in larger models like ESM2-3B, which cannot be fully fine-tuned on local GPUs due to its size but can be efficiently trained using LoRA with rank 4, demonstrating LoRA’s practical effectiveness for large pLMs. This highlights LoRA’s ability to deliver competitive performance with significantly reduced computational resources compared to full fine-tuning. These results demonstrate the effectiveness of the full and PEFT fine-tuning over the pre-trained models and the machine learning techniques, reflected when compared to the existing methods on both independent datasets (TF and TFPM independent sets). Full fine-tuning of ProtBert using transcription data failed to achieve good results primarily due to optimization difficulties during training. Such failures often stem from vanishing gradients in early training stages, where the model’s parameter updates become very small and the training does not progress effectively. This issue leads to poor convergence and performance stagnation on the new task.

Our analysis using the fully fine-tuned ESM2-650M model demonstrates that the attention mechanisms/weights captured by the different heads and layers effectively highlight biologically meaningful amino acid fragments, particularly those corresponding to experimentally validated motifs such as “ERKRRRNR.” By comparing attention patterns with STREME-derived motifs, we show that full fine-tuning enhances the model’s ability to capture functional interactions within transcription factor sequences. The complementary behaviors of different attention heads and layers, with one emphasizing motif-level relationships and another reinforcing local structural consistency, provide interpretable insights into the model’s decision-making process. These findings support the reliability of the proposed framework and underscore the value of attention-based interpretation in protein sequence classification tasks.

In the future work, we could involve using larger and more advanced models, like the recent ESM-2 variants, as they have been shown to improve performance in protein-related tasks. Additionally, integrating retrieval-based methods like RAG-ESM or leveraging enhanced structural and functional annotations may further improve model interpretability and accuracy. Addressing the limitations of resource constraints and model complexity, such as developing more efficient or resource-friendly versions, is also a promising direction for future research.

## Data availability

The data, source code, and trained models are made available at GitHub at https://github.com/mirtanveer/Fine_Tune_TF. The full and PEFT fine-tuning of the ESM2 models and the inference/prediction on the TF independent dataset using the LoRA-ESM2-650M trained model can be carried out using the Google Colab notebooks attached to the GitHub https://github.com/MirTanveer/Fine-Tune-TF/tree/main/Notebooks.

## Supplementary Data

Some partial experimental results are described in the supplementary material.

