## Supplementary File for "Fine-Tuning Protein Language Models Enhances the Identification and Interpretation of the Transcription Factors"

Table S1: Cross-validation performance of the machine learning models for the TF identification using PSSM features

| Encoding | Model | Acc | Sn | Sp | MCC | F1_score | AUC |
| --- | --- | --- | --- | --- | --- | --- | --- |
| AAC­_PSSM | RF | 0.844 | 0.861 | 0.829 | 0.692 | 0.872 | 0.923 |
|  | ETC | 0.853 | 0.882 | 0.824 | 0.706 | 0.856 | 0.935 |
|  | XGB | 0.809 | 0.838 | 0.778 | 0.620 | 0.797 | 0.876 |
|  | LGBM | 0.840 | 0.865 | 0.814 | 0.680 | 0.830 | 0.913 |
|  | CatBoost | 0.789 | 0.762 | 0.817 | 0.579 | 0.781 | 0.873 |
| AB_PSSM | RF | 0.854 | 0.858 | 0.85 | 0.708 | 0.868 | 0.929 |
|  | ETC | 0.874 | 0.838 | 0.911 | 0.752 | 0.861 | 0.939 |
|  | XGB | 0.806 | 0.735 | 0.778 | 0.610 | 0.806 | 0.881 |
|  | LGBM | 0.866 | 0.865 | 0.860 | 0.720 | 0.859 | 0.912 |
|  | CatBoost | 0.858 | 0.838 | 0.877 | 0.718 | 0.843 | 0.927 |
| Tri_gram_PSSM | RF | 0.884 | 0.843 | 0.925 | 0.774 | 0.875 | 0.934 |
|  | ETC | 0.899 | 0.831 | 0.966 | 0.805 | 0.894 | 0.932 |
|  | XGB | 0.802 | 0.793 | 0.807 | 0.602 | 0.834 | 0.882 |
|  | LGBM | 0.863 | 0.860 | 0.860 | 0.720 | 0.859 | 0.911 |
|  | CatBoost | 0.789 | 0.761 | 0.817 | 0.579 | 0.781 | 0.873 |
| DP_PSSM | RF | 0.875 | 0.862 | 0.889 | 0.750 | 0.862 | 0.931 |
|  | ETC | 0.871 | 0.858 | 0.884 | 0.743 | 0.874 | 0.935 |
|  | XGB | 0.871 | 0.855 | 0.888 | 0.743 | 0.859 | 0.936 |
|  | LGBM | 0.862 | 0.858 | 0.868 | 0.726 | 0.862 | 0.938 |
|  | CatBoost | 0.867 | 0.854 | 0.880 | 0.735 | 0.868 | 0.943 |
| Smoothed_PSSM | RF | 0.786 | 0.771 | 0.831 | 0.580 | 0.770 | 0.862 |
|  | ETC | 0.770 | 0.795 | 0.742 | 0.541 | 0.764 | 0.851 |
|  | XGB | 0.654 | 0.682 | 0.628 | 0.307 | 0.698 | 0.710 |
|  | LGBM | 0.784 | 0.795 | 0.775 | 0.573 | 0.768 | 0.861 |
|  | CatBoost | 0.771 | 0.769 | 0.775 | 0.543 | 0.759 | 0.859 |
| *K_*separated_bigram_PSSM | RF | 0.871 | 0.829 | 0.913 | 0.747 | 0.873 | 0.931 |
|  | ETC | 0.888 | 0.853 | 0.923 | 0.777 | 0.879 | 0.936 |
|  | XGB | 0.847 | 0.834 | 0.868 | 0.696 | 0.844 | 0.913 |
|  | LGBM | 0.865 | 0.850 | 0.880 | 0.730 | 0.863 | 0.921 |
|  | CatBoost | 0.875 | 0.860 | 0.889 | 0.749 | 0.880 | 0.936 |
| PSe_PSSM | RF | 0.866 | 0.879 | 0.854 | 0.734 | 0.869 | 0.914 |
|  | ETC | 0.877 | 0.887 | 0.868 | 0.756 | 0.879 | 0.943 |
|  | XGB | 0.844 | 0.853 | 0.837 | 0.691 | 0.846 | 0.902 |
|  | LGBM | 0.861 | 0.874 | 0.848 | 0.724 | 0.863 | 0.919 |
|  | CatBoost | 0.869 | 0.879 | 0.859 | 0.737 | 0.867 | 0.943 |

Table S2: Independent performance of the machine learning models for the TF identification using PSSM features

| Encoding | Model | Acc | Sn | Sp | MCC | F1_score | AUC |
| --- | --- | --- | --- | --- | --- | --- | --- |
| AAC_PSSM | RF | 0.849 | 0.867 | 0.83 | 0.698 | 0.851 | 0.927 |
|  | ETC | 0.867 | 0.924 | 0.811 | 0.740 | 0.875 | 0.931 |
|  | XGB | 0.834 | 0.896 | 0.773 | 0.674 | 0.844 | 0.919 |
|  | LGBM | 0.872 | 0.886 | 0.858 | 0.745 | 0.874 | 0.934 |
|  | CatBoost | 0.793 | 0.767 | 0.821 | 0.588 | 0.788 | 0.881 |
| AB_PSSM | RF | 0.825 | 0.849 | 0.802 | 0.651 | 0.829 | 0.902 |
|  | ETC | 0.845 | 0.811 | 0.877 | 0.690 | 0.839 | 0.914 |
|  | XGB | 0.765 | 0.821 | 0.707 | 0.531 | 0.776 | 0.844 |
|  | LGBM | 0.806 | 0.83 | 0.784 | 0.613 | 0.811 | 0.883 |
|  | CatBoost | 0.793 | 0.801 | 0.784 | 0.585 | 0.794 | 0.883 |
| Tri_gram_PSSM | RF | 0.872 | 0.840 | 0.906 | 0.746 | 0.868 | 0.937 |
|  | ETC | 0.892 | 0.850 | 0.933 | 0.785 | 0.886 | 0.942 |
|  | XGB | 0.816 | 0.838 | 0.801 | 0.632 | 0.818 | 0.882 |
|  | LGBM | 0.807 | 0.839 | 0.784 | 0.613 | 0.811 | 0.883 |
|  | CatBoost | 0.794 | 0.767 | 0.821 | 0.585 | 0.788 | 0.882 |
| DP_PSSM | RF | 0.863 | 0.849 | 0.877 | 0.726 | 0.861 | 0.915 |
|  | ETC | 0.891 | 0.886 | 0.897 | 0.783 | 0.890 | 0.929 |
|  | XGB | 0.873 | 0.887 | 0.858 | 0.745 | 0.874 | 0.923 |
|  | LGBM | 0.868 | 0.868 | 0.867 | 0.735 | 0.867 | 0.927 |
|  | CatBoost | 0.872 | 0.868 | 0.878 | 0.745 | 0.872 | 0.928 |
| Smoothed_PSSM | RF | 0.740 | 0.716 | 0.764 | 0.481 | 0.734 | 0.842 |
|  | ETC | 0.731 | 0.716 | 0.745 | 0.462 | 0.727 | 0.824 |
|  | XGB | 0.653 | 0.681 | 0.627 | 0.307 | 0.698 | 0.710 |
|  | LGBM | 0.758 | 0.754 | 0.746 | 0.500 | 0.751 | 0.857 |
|  | CatBoost | 0.731 | 0.731 | 0.727 | 0.462 | 0.732 | 0.831 |
| *K_*separated_bigram_PSSM | RF | 0.853 | 0.83 | 0.877 | 0.708 | 0.850 | 0.932 |
|  | ETC | 0.896 | 0.878 | 0.916 | 0.793 | 0.894 | 0.952 |
|  | XGB | 0.839 | 0.867 | 0.813 | 0.680 | 0.844 | 0.926 |
|  | LGBM | 0.897 | 0.924 | 0.868 | 0.793 | 0.899 | 0.946 |
|  | CatBoost | 0.905 | 0.906 | 0.896 | 0.801 | 0.904 | 0.963 |
| PSe_PSSM | RF | 0.831 | 0.849 | 0.812 | 0.660 | 0.833 | 0.925 |
|  | ETC | 0.858 | 0.877 | 0.839 | 0.717 | 0.861 | 0.936 |
|  | XGB | 0.806 | 0.802 | 0.811 | 0.613 | 0.805 | 0.888 |
|  | LGBM | 0.845 | 0.858 | 0.831 | 0.688 | 0.930 | 0.933 |
|  | CatBoost | 0.868 | 0.854 | 0.887 | 0.736 | 0.870 | 0.934 |

Table S3: The runtime comparison between the fully fine-tuned and LoRA models. The runtime values are described in seconds.

| Runtime (*seconds*)/Models | ESM2-8M | ESM2-35M | ESM2-150M | ESM2-650M |
| --- | --- | --- | --- | --- |
| Fully fine-tuned models | 245.013 | 508.849 | 2256.939 | 36972.313 |
| LoRA models | 172.641 | 362.472 | 1250.000 | 12350.768 |

Table S4: The performance of the full fine-tuned ESM2 models on the TFPM independent dataset for the prediction of TF interactions with methylated DNA.

| Model | Acc | Sn | Sp | MCC | F1_score | AUC |
| --- | --- | --- | --- | --- | --- | --- |
| ESM2-8M | 0.593 ± 0.043 | 0.414 ± 0.047 | 0.726 ± 0.044 | 0.149 ± 0.067 | 0.511 ± 0.042 | 0.490 ± 0.045 |
| ESM2-35M | 0.589 ± 0.033 | 0.453 ± 0.047 | 0.725 ± 0.043 | 0.186 ± 0.067 | 0.492 ± 0.042 | 0.524 ± 0.044 |
| ESM2-150M | 0.429 ± 0.048 | 0.368 ± 0.058 | 0.542 ± 0.084 | -0.087 ± 0.099 | 0.437 ± 0.058 | 0.452 ± 0.058 |
| ESM2-650M | 0.429 ± 0.049 | 0.338 ± 0.057 | 0.596 ± 0.083 | -0.064 ± 0.101 | 0.429 ± 0.060 | 0.432 ± 0.060 |

Table S5: The performance of the ESM2 models on the TFPM independent dataset for the prediction of TF interactions with methylated DNA using LoRA.

| Model | Acc | Sn | Sp | MCC | F1_score | AUC |
| --- | --- | --- | --- | --- | --- | --- |
| LoRA-ESM2-8M | 0.467 ± 0.049 | 0.295 ± 0.054 | 0.784 ± 0.070 | 0.085 ± 0.097 | 0.543 ± 0.058 | 0.415 ± 0.062 |
| LoRA-ESM2-35M | 0.495 ± 0.046 | 0.352 ± 0.055 | 0.756 ± 0.067 | 0.111 ± 0.089 | 0.543 ± 0.058 | 0.472 ± 0.058 |
| LoRA-ESM2-150M | 0.619 ± 0.046 | 0.943 ± 0.028 | 0.027 ± 0.028 | -0.064 ± 0.089 | 0.525 ± 0.057 | 0.761 ± 0.035 |
| LoRA-ESM2-650M | 0.475 ± 0.045 | 0.322 ± 0.055 | 0.756 ± 0.069 | 0.082 ± 0.089 | 0.531 ± 0.058 | 0.440 ± 0.060 |
